# drICE restrains DIAP2-mediated inflammatory signalling and intestinal inflammation

**DOI:** 10.1101/2020.05.29.122978

**Authors:** Christa Kietz, Vilma Pollari, Ida-Emma Tuominen, Paulo S Ribeiro, Pascal Meier, Annika Meinander

## Abstract

The *Drosophila* IAP protein, DIAP2, is a key mediator of NF-κB signalling and is required for immune activation, both locally in the intestinal epithelia and during systemic, fat body-induced responses. We have found that transgenic expression of DIAP2 induces inflammation in the intestine, but not in the fat body, indicating a need for regulating DIAP2 in microbiotic environments. We describe the *Drosophila* caspase drICE, a known interaction partner of DIAP2, as a regulator of DIAP2 and NF-κB signalling in the intestinal epithelium. drICE acts by cleaving, and interfering with DIAP2’s ability to ubiquitinate and, thereby, activate mediators of the NF-κB pathway. The drICE-cleaved form of DIAP2 is, moreover, unable to induce inflammation during basal conditions in the intestine. Interestingly, cleavage of DIAP2 does not interfere with pathogen-induced signalling, suggesting that drICE protects from immune responses induced by resident microbes. Accordingly, the negative regulatory effect of drICE is lost when rearing flies axenic. Hence, we show that drICE halts unwanted inflammatory signalling in the intestine by forming an inhibitory complex with DIAP2, interfering with the ability of DIAP2 to induce downstream signalling and NF-κB target gene activation.

## Introduction

Innate immune responses are initiated by pattern recognition receptors (PRRs) that recognize pathogen-associated molecular patterns (PAMPs), and danger-associated molecular patterns (DAMPs). Activation of PRRs leads to the induction of microbicidal and pro-inflammatory responses, and culminates in elimination of the activating molecule and subsequent return to cellular homeostasis (Takeuchi & Akira, 2010). Proper regulation of inflammatory signalling is crucial, as de-regulation at any step can be detrimental for the organism. One of the key players in the inflammatory response is the NF-κB family of transcription factors that regulate the expression of numerous inflammatory genes. Constitutively active NF-κB signalling is characteristic of chronic inflammations and increased NF-κB activity has been connected to irritable bowel diseases, such as ulcerative colitis and Crohn’s disease (Atreya et al., 2008, Viennois et al., 2013).

Inhibitor of apoptosis proteins (IAPs) influence ubiquitin-dependent pathways that modulate innate immune signalling by activation by NF-κB (Darding & Meier, 2012). IAPs were first identified in insect baculoviruses as potent inhibitors of cell death (Crook et al., 1994, Birnbaum et al., 1994) and have subsequently been identified in both vertebrates and invertebrates. Cellular and viral IAPs are characterized by the presence of one or more caspase-binding baculoviral repeat (BIR) domains that are essential for their anti-apoptotic properties (Birnbaum et al., 1994, Hinds et al., 1999), as well as a RING (really interesting new gene) domain, providing them with E3 ligase activity (Vaux and Silke, 2005). *Drosophila* carries two *bona fide* IAP genes, *Drosophila* inhibitor of apoptosis protein (DIAP) 1 and 2 (Hay et al., 1995). DIAP1 functions mainly as a suppressor of cell death, whereas DIAP2, although also able to decrease the apoptotic threshold of the cell, has its main function in inflammatory signalling (Kleino et al., 2005, Gesellchen et al., 2005, Leulier et al., 2006a, Huh et al., 2007). The most extensively studied mammalian IAPs, i.e. cellular IAP1/2 and XIAP (X-chromosome-linked IAP), and *Drosophila* DIAP2 harbour furthermore, an UBA domain (ubiquitin-associated domain) enabling them to interact with poly-ubiquitin chains (Gyrd-Hansen et al., 2008, Blankenship et al., 2009).

In contrast to mammals, *Drosophila* relies solely on an innate immune defence when combating pathogenic infections. One of the key components of the fly’s immune system is the production and secretion of antimicrobial peptides (AMPs). The production is regulated by two NF-κB pathways, namely, the Toll pathway and the IMD (immune deficient) pathway (Hetru & Hoffman, 2009). Upon a Gram-negative or Gram-positive systemic infection, the IMD and Toll pathways, respectively, activate the production of AMPs in the fat body (Lemaitre & Hoffman, 2007, Kurata, 2010). However, during local immune responses in the gut, the IMD pathway solely controls the generation of AMPs (Ferrandon et al., 1998, Tzou et al., 2000). The IMD pathway is activated by PGRP-LCx receptors recognizing diaminopimelate (DAP) –type peptidoglycans (PGN), which are components of the cell wall of Gram-negative bacteria (Choe et al., 2002, Gottar et al., 2002, Leulier et al., 2003). Activation of PGRP-LCx leads to the recruitment of the adaptor proteins IMD and dFADD, and the initiator caspase DREDD (Choe et al., 2005). DREDD-mediated cleavage of IMD exposes a conserved IAP-binding motif that recruits DIAP2 to the complex, stimulating DIAP2-mediated K63-ubiquitination of IMD, DREDD and the IKK Kenny (Paquette et al., 2010, Meinander et al., 2012, Aalto et al., 2019). Ubiquitination of DREDD is needed for cleavage and nuclear localization of the NF-κB protein Relish (Stoven et al., 2003, Meinander et al., 2012), whereas ubiquitination of IMD has been suggested to recruit the *Drosophila* mitogen-activated protein kinase kinase kinase dTAK1 and the Relish kinase complex IRD5/Kenny to the IMD signalling complex (Ferrandon et al., 2007).

Vertebrate and invertebrate organisms are in continuous contact with a diverse array of resident microorganisms (Qin et al., 2010). One key interface for host-microbe interactions, in both humans and *Drosophila*, is the epithelial layer of the gut (Qin et al., 2010, Apidianakis and Rahme, 2011). In addition of being a platform for beneficial host-microbe interactions, the gut epithelium serves as the first line of defence against pathogens entering the body. In order for the organism to mount an efficient immune response against pathogenic bacteria, and simultaneously allowing commensal bacteria to interact with the host, inflammatory signalling needs to be carefully regulated. Here, we report a novel role of the *Drosophila* caspase-3 homologue drICE as a negative regulator of the IMD pathway in the intestinal epithelium. Our results show that drICE restrains inflammatory signalling induced by commensal bacteria in fly intestine by interacting with DIAP2, interfering with its E3 ligase activity. We also show that transgenic expression of DIAP2 leads to chronic inflammation only in the presence of commensal bacteria, which elucidates the need for DIAP2 to be regulated in microbiotic environments.

## Results

### DIAP2 induces chronic inflammation in the fly intestine

The *Drosophila* IAP protein DIAP2 is required for inducing the Relish-dependent IMD pathway and mounting immune responses upon Gram-negative infection, both locally in the epithelial layers of the gut and trachea, and upon systemic infections in the cells of the fat body (Kleino et al., 2005, Leulier et al., 2006a). However, it is still unknown how the activity of DIAP2 is regulated during basal conditions and upon pathogen-induced infections. To investigate how DIAP2-mediated inflammatory responses are regulated, we first tested the impact of DIAP2 on inflammatory NF-κB activation. For this purpose, we made transgenic flies, in which expression of DIAP2 was induced via the UAS-Gal4 system (**Figure 1A)**. Analysis of the expression of the Relish target genes *Drosocin* and *Diptericin* in whole-fly lysates showed that these AMPs are induced in the DIAP2-expressing flies compared to *CantonS* and *Ubiquitin-Gal4* (*UbiGal4*) control flies (**Figure 1B**). To investigate the origin of the DIAP2-induced inflammation, we used the *Diptericin-LacZ* reporter system to compare *Diptericin* expression in two of the major immune organs of the fly, the gut and the fat body (Lemaitre and Hoffmann, 2007). We found that transgenic expression of DIAP2 leads to a spontaneous induction of *Diptericin* in the *Drosophila* midgut, but not in the fat body (**Figure 1C, D)**. As DIAP2 overexpression did not induce an inflammatory response in the fat body, we compared the endogenous expression of DIAP2 in the intestine and the fat body of *CantonS* wild type flies. Interestingly, while all endogenous DIAP2 was cleaved during basal conditions in the intestine, DIAP2 was not cleaved in the fat body (**Figure 1E**). When examining the protein levels of DIAP2 in DIAP2-expressing flies, we found that full-length DIAP2 could be detected in the intestines of these flies, indicating that an increased expression of full length DIAP2 (**Figure 1F**) induces inflammation only in the intestine.

**Figure 1.**
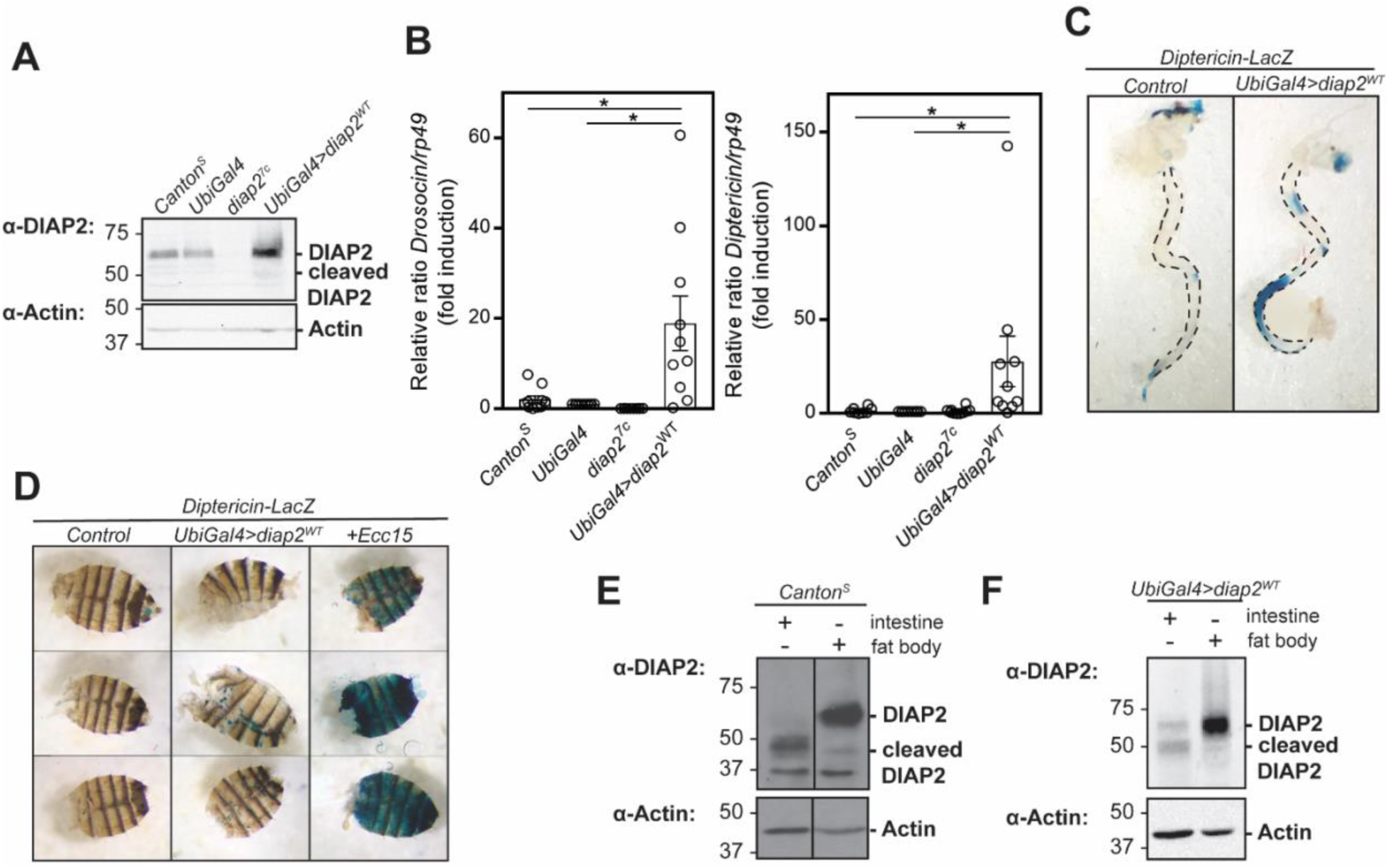
DIAP2 induces chronic inflammation in the fly intestine. A) Whole fly lysates from *CantonS, UbiGal4, diap2*_*7c*_ and *UbiGal4;UAS-diap2*_*WT*_ flies were analysed by Western blotting with α-DIAP2 and α-Actin antibodies, n=3. B) Relative *Droso*c*in* and *Diptericin* mRNA levels analysed with qRT-PCR in adult *CantonS, UbiGal4, diap2*_*7c*_ and *UbiGal4;UAS-diap2*_*WT*_ flies. Error bars indicate ±SEM from ten independent experimental repeats, * p < 0.05. C, D) Adult female intestines (C) and abdomens (D) from *Diptericin-LacZ* and *UbiGal4;UAS-diap2*_*WT*_*/Diptericin-LacZ* flies were dissected and stained for β-galactosidase activity, n=3. The last lane in panel D shows a positive control for fat body activation induced by septic infection with *Ecc15*. E, F) Intestines and fat bodies from adult female *CantonS* (E) or *UbiGal4;UAS-diap2*_*WT*_ (F) flies were dissected and lysed, and analysed by Western blotting with α-DIAP2 and α-Actin antibodies, n=3.

### drICE restrains intestinal activation of the IMD pathway and gut hyperplasia

Caspases belong to a family of conserved cysteine-dependent endoproteases that cleave their substrates after specific aspartic residues (Van Opdenbosch and Lamkanfi, 2019). The *Drosophila* effector caspase drICE is one of the key inducers of apoptosis in the fly (Fraser and Evan, 1997). DIAP2 and drICE have been shown to interact and form a stable complex, wherein DIAP2 inhibits the activity of drICE by binding to, and ubiquitinating the caspase (Ribeiro et al., 2007). As we found DIAP2 to be constitutively cleaved in the intestine, we wanted to further investigate a possible role for drICE in the regulation of DIAP2-mediated inflammatory signalling. For this purpose, we measured the expression of NF-κB target genes in *drICE*_*17*_ mutant flies and in *drICE RNAi* flies. *drICE*_*17*_ flies encode a highly unstable form of the drICE protein, reported to express less than 5% of the wild type-expressed drICE (Xu et al., 2006), and neither *drICE*_*17*_ nor *drICE RNAi* flies have detectable expression of drICE (**Figure 2A**). We detected a significantly higher expression of the Relish target genes *Drosocin* and *Diptericin* during basal conditions in whole fly lysates from both *drICE*_*17*_ (**Figure 2B**) and *drICE RNAi* flies (**Figure 2C**) and, agreeingly, a lower basal *Drosocin* and *Diptericin* expression in flies overexpressing drICE (**Figure 2D**), suggesting that drICE acts as a negative regulator of IMD signalling.

**Figure 2.**
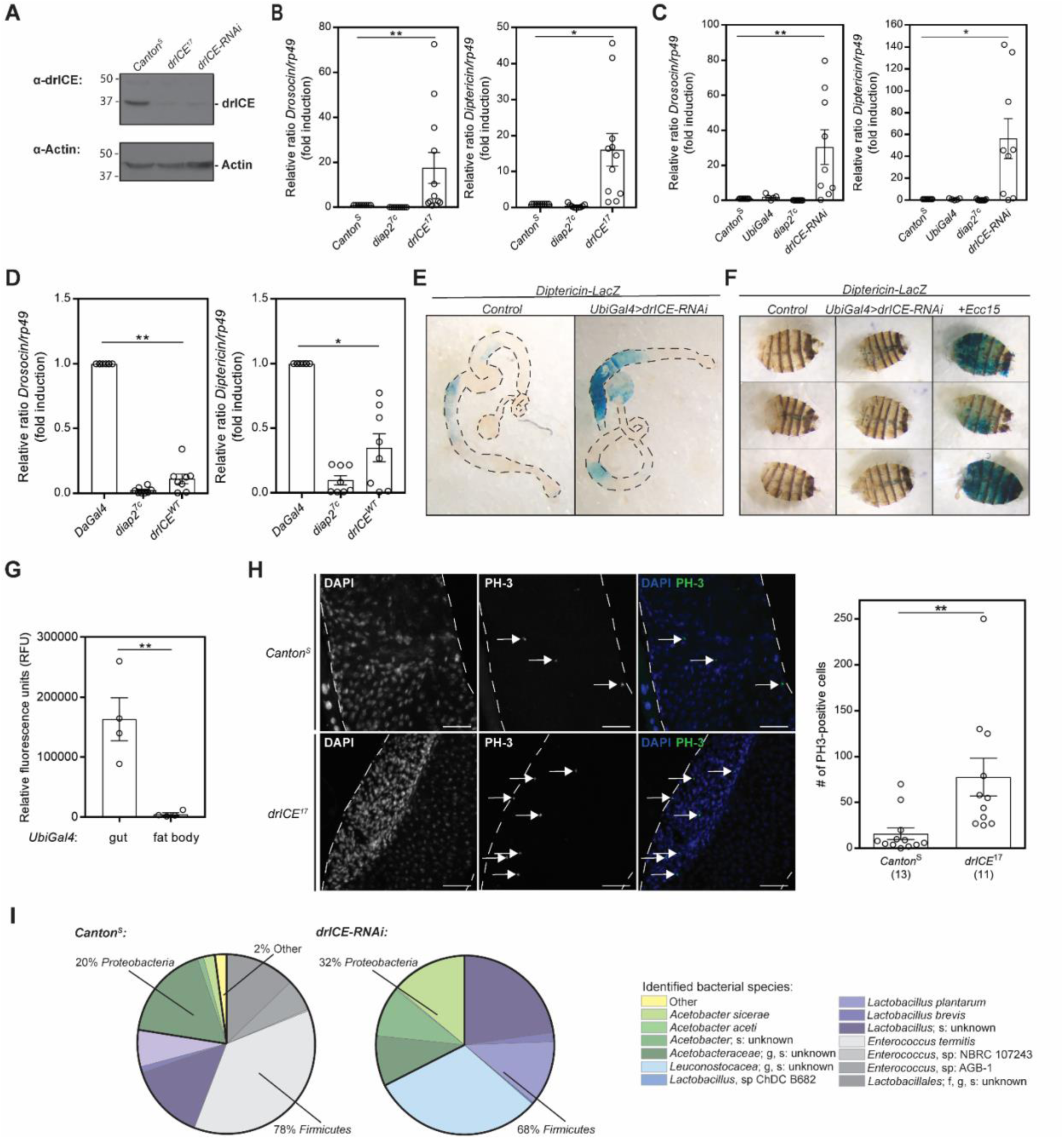
drICE restrains intestinal activation of the IMD pathway and gut hyperplasia. A) Whole fly lysates from *CantonS, drICE*_*17*_ and *UbiGal4;UAS-drICE-RNAi* flies were analysed by Western blotting with α-drICE and α-Actin antibodies, n=3. B, C, D) Relative *Drosocin* and *Diptericin* mRNA levels analysed with qRT-PCR in adult *CantonS, diap2*_*7c*_ and *drICE*_*17*_ flies (B), in *CantonS, UbiGal4, diap2*_*7c*_, and *UbiGal4;UAS-drICE-RNAi* (C) flies or in *DaGal4, diap2*_*7c*_ and *UAS-drICE*_*WT*_;*DaGal4* flies. Error bars indicate ±SEM from at least eight independent experimental repeats, * p < 0.05, ** p < 0.01. E, F) Adult female intestines (E) and abdomens (F) from *Diptericin-LacZ* and *UbiGal4;UAS-drICE-RNAi/Diptericin-LacZ* flies were dissected and stained for β-galactosidase activity, n=3. The last lane in panel E shows a positive control for fat body activation induced by septic infection with *Ecc15*. G) Adult female intestines or fat bodies from *UbiGal4* flies were dissected and lysed, and the caspase-3/-7 activity was assessed after addition of Apo-ONE reagent by measuring fluorescence at 499/521 nm. Error bars indicate ± SEM from four independent experiments, ** p < 0.01. H) Intestines from adult *Canton*_*S*_ and *drICE*_*17*_ flies were dissected and stained for phosphohistone H3 (PH-3) (green) and DAPI (blue). The PH3-positive cells are marked with arrows and the scale bar indicates 50 µm. All PH3-positive cells in midguts prepared and stained were counted for statistics, error bars indicate ±SEM from three independent experimental repeats, ** p < 0.01. The number of intestines analysed is indicated in brackets. I) Bacterial 16S rRNA metagenomics analysis of the 1V-3V region in *CantonS* and *drICE-RNAi* flies. Colours indicate identified operational taxonomic units (OTUs). Black lines indicate proportions of *Proteobacteria* and *Firmicutes*, n=1.

To investigate if drICE-mediated regulation of the IMD pathway is tissue-specific, we examined the *Diptericin* expression in the gut and fat body of *drICE RNAi* flies carrying the *Diptericin-LacZ* reporter gene. Similarly, as DIAP2 overexpression only induced inflammation in the intestine, loss of *drICE* induced *Diptericin* expression in the midgut, but not in the fat body (**Figure 2E, F)**. We, furthermore, found that active drICE, assessed by measuring DEVD-activity, was readily detected in dissected fly guts but not in the fat body (**Figure 2G**), supporting the notion of a tissue-specific role of drICE. Intestinal inflammation is associated with midgut hyperplasia and proliferating cells can be detected in *Drosophila* by staining the proliferation marker phospho-histone H3 (PH-3) (Amcheslavsky et al., 2009). We found *drICE*_*17*_ mutant flies to have a significantly increased number of proliferating cells in the midgut, compared to *CantonS* control flies (**Figure 2H**). Another factor associated with chronic inflammation is dysbiosis of the gut microbiome. When profiling the bacterial composition of wild type and *drICE RNAi* flies by 16S sequencing, we found the *drICE RNAi* flies to have a higher ratio of *Proteobacteria* to *Firmicutes* compared to wild-type *CantonS* flies (**Figure 2I**), a notion that has been associated with increased gut inflammation in both flies and humans (Clark et al., 2015, Clemente et al., 2012). Taken together, our results show that drICE is a negative regulator of IMD signalling, needed for maintaining gut homeostasis.

### The catalytic activity of drICE is needed to regulate the IMD pathway in the intestine

To investigate whether the activity of DIAP2 is restrained by drICE-mediated cleavage, we generated transgenic flies expressing drICE_WT_ or the catalytically inactive drICE_C211A_ mutant. While *drICE RNAi* eliminated the endogenous drICE, the transgenic expressed drICE remained high in a drICE RNAi background (**Figure 3A**). When we analysed the expression of the Relish target genes *Drosocin* and *Diptericin* in whole-fly lysates during basal conditions, we found that only drICE_WT_, but not drICE_C211A_ is able to restrain the AMP expression induced by loss of *drICE* (**Figure 3B**). As drICE cleaves DIAP2 at D100 (Ribeiro et al., 2007), we also made transgenic flies expressing the drICE-cleaved DIAP2_Δ100_, consisting of the 101 to 338 last amino acids of DIAP2, under control of the UAS-Gal4 system. When expressing DIAP2_Δ100_ or DIAP2_WT_ in a *diap2*_*7c*_ mutant background, only the full-length DIAP2 was able to induce expression of *Drosocin* and *Diptericin* during basal conditions (**Figure 3C**). Similarly, we found expression of DIAP2_WT_, but not DIAP2_Δ100_ to induce local *Diptericin* expression in the midgut (**Figure 3D**).

**Figure 3.**
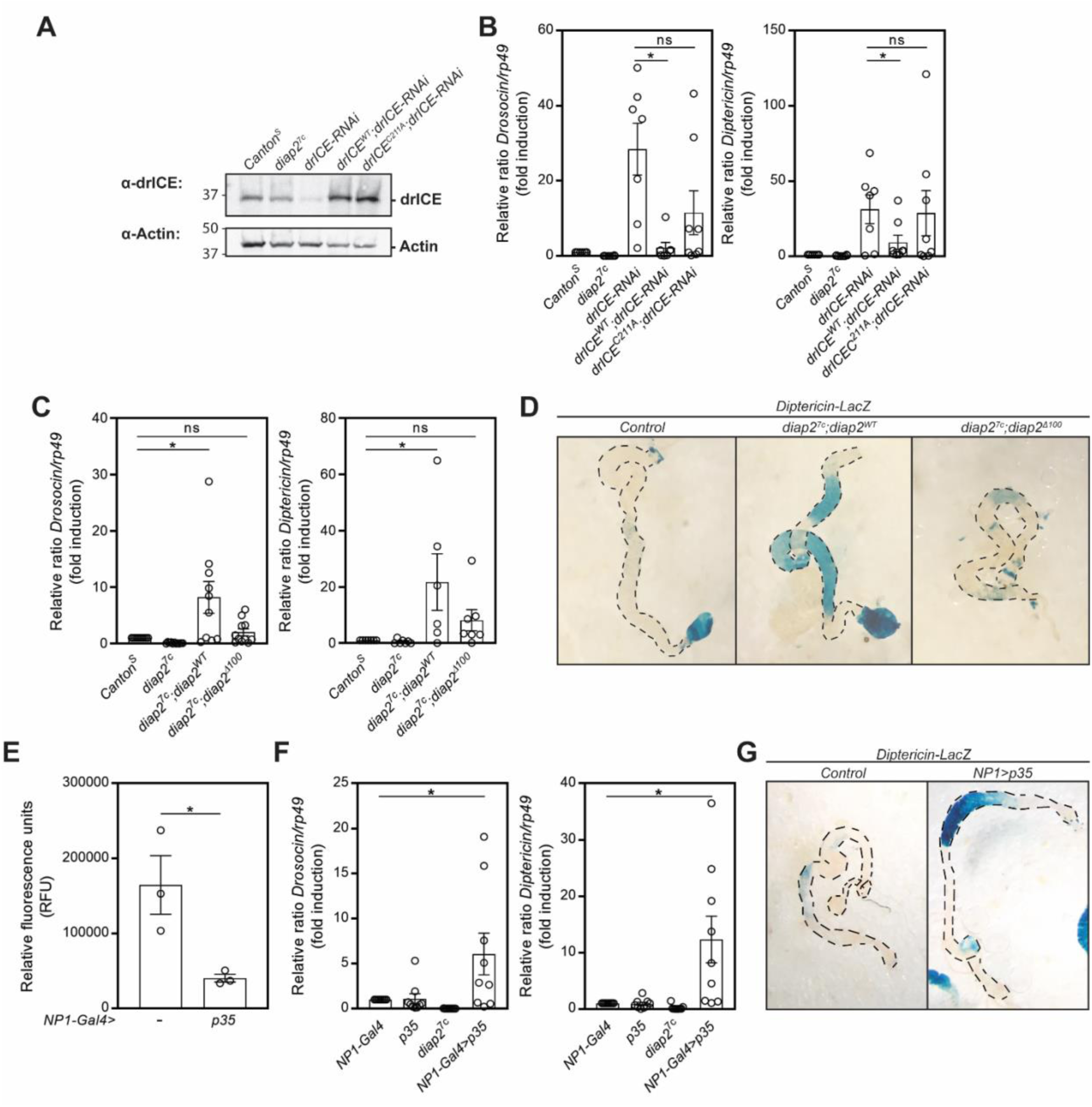
The catalytic activity of drICE is needed to regulate the IMD pathway in the intestine. A) Whole fly lysates from *CantonS, diap2*_*7c*_, *UbiGal4*;*UAS*-*drICE-RNAi, UAS*-*drICE*_*WT*_*/UbiGal4;UAS-drICE-RNAi* and *UAS*-*drICE*_*C211A*_*/UbiGal4;UAS-drICE-RNAi* were analysed by Western blotting with α-drICE and α-Actin antibodies, n=3. B) Relative *Drosocin* and *Diptericin* mRNA levels analysed with qRT-PCR in adult *CantonS, diap2*_*7c*_, *UbiGal4*;*UAS*-*drICE-RNAi, UAS*-*drICE*_*WT*_*/UbiGal4;UAS-drICE-RNAi* and *UAS*-*drICE*_*C211A*_*/UbiGal4;UAS-drICE-RNAi* flies. Error bars indicate ±SEM from seven independent experimental repeats. C) Relative *Drosocin* and *Diptericin* mRNA levels analysed with qRT-PCR in adult *CantonS, diap2*_*7c*_, *diap2*_*7c*_;*UAS-diap2*_*WT*_*/DaGal4* and in *diap2*_*7c*_;*UAS-diap2*_*Δ100*_*/DaGal4* flies. Error bars indicate ±SEM from at least six independent experimental repeats, ns=non-significant, * p < 0.05, ** p < 0.01. D) Adult female intestines from *DaGal4, Diptericin-LacZ, diap2*_*7c*_;*UAS-diap2*_*WT*_*/DaGal4, Diptericin-LacZ* and *diap2*_*7c*_;*UAS*-*diap2*_*Δ100*_*/DaGal4, Diptericin-LacZ* flies were dissected and stained for β-galactosidase activity, n=3. E) Adult female intestines from *NP1-Gal4* and *NP1-Gal4;UAS-p35* flies were dissected and lysed, and the caspase-3/-7 activity was assessed after addition of Apo-ONE reagent by measuring fluorescence at 499/521 nm. Error bars indicate ±SEM from three independent experiments, * p < 0.05. F) Relative *Drosocin* and *Diptericin* mRNA levels analysed with qRT-PCR in adult *NP1-Gal4, UAS-p35, diap2*_*7c*_ and *NP1-Gal4;UAS-p35* flies. Error bars indicate ±SEM from ten independent experimental repeats, * p < 0.05. G) Adult female intestines from *DaGal4, Diptericin-LacZ* and *NP1-Gal4;UAS-p35/Diptericin-LacZ* flies were dissected and stained for β-galactosidase activity, n=3.

To verify that drICE is the caspase responsible for cleaving DIAP2, we expressed the effector caspase inhibitor p35 in the intestinal epithelial cells with the enterocyte-specific driver *NP1-Gal4*. When cleaved by a caspase, the viral caspase inhibitor p35 acts as a suicide substrate by trapping the catalytic machinery of the caspase via a covalent thioacyl linkage (Stennicke & Salvesen, 2006). p35 has been shown to inhibit the *Drosophila* effector caspases drICE and DCP-1 (Kim et al., 2014), which both recognise the DEVD amino acid sequence (Fraiser et al., 1997, Song et al., 1997). However, drICE has been shown to be the only *Drosophila* effector caspase to interact with DIAP2 (Leulier et al., 2006b). We found that expression of p35 in intestinal enterocytes led to reduced effector caspase activity **(Figure 3E)** and increased expression of the Relish target genes *Drosocin* and *Diptericin* (**Figure 3F**). Likewise, a local induction in *Diptericin* expression in the intestine was detected in *Diptericin-LacZ* flies expressing the p35 caspase inhibitor (**Figure 3G**). Our results indicate, thus, that the catalytic cysteine and, thereby, the covalent bond formed between DIAP2 and drICE is needed for drICE to regulate the activity of the IMD pathway, and that p35, by trapping the catalytic cysteine of drICE, interferes with the ability of drICE to restrain IMD signalling. Although the initiator caspases DREDD and DRONC have been shown to interact with DIAP2 (Quinn et al., 2000 Guntermann et al., 2014), neither DREDD nor DRONC are inhibited by p35 (Meier et al., 2000, Kim et al, 2014), suggesting that drICE alone inhibits DIAP2-mediated activation of Relish target genes by its catalytic activity.

### drICE interferes with the with the E3 ligase activity of DIAP2

To investigate how drICE-mediated cleavage of DIAP2 regulates IMD signalling, we examined the ability of DIAP2 to ubiquitinate its substrates in the presence or absence of active drICE. DIAP2-mediated K63-linked ubiquitination of the IMD pathway components IMD and DREDD has been shown to be required for IMD signalling, and the *Drosophila* NEMO, IKKγ or Kenny, has recently been identified as a target of K63-linked ubiquitination mediated by DIAP2 (Paquette et al., 2010, Meinander et al., 2012, Aalto et al., 2019). While drICE activity was induced by overexpressing drICE in S2 cells, drICE was inhibited by overexpression of the catalytically inactive drICE_C211A_ or by treatment with synthetic caspase-3 inhibitor Z-DEVD-FMK. Inhibition of caspase activity was verified by analysing DEVD-activity and drICE cleavage (**Figure 4A, B**). To investigate whether drICE interferes with the E3-ligase activity of DIAP2, we co-transfected S2-cells with DIAP2_WT_ and DREDD-V5 or Kenny-V5. Ubiquitin chains were pulled down with recombinant GST-TUBE from cell lysates under denaturing conditions. When overexpressing drICE_WT_ we, indeed, detected a decreased amount of ubiquitin chains bound to DREDD and Kenny (**Figure 4C, E, lanes 3, and 4D, F**). Agreeingly, a stronger high molecular smear of DREDD and Kenny was detected after treatment with Z-DEVD-FMK and when overexpressing drICE_C211A_ (**Figure 4C, E, lanes 4 and 5, and 4D, F**). Likewise, we found a reduced ability of the drICE-cleaved DIAP2_Δ100_ to ubiquitinate DREDD compared to DIAP2_WT_. However, some ubiquitination can still be induced by DIAP2_Δ100_ compared to the DIAP2 RING mutant DIAP2_F462A_ (**Figure 4G, H**). Taken together these results indicate that drICE restrains the ability of DIAP2 to ubiquitinate the IMD pathway inducers DREDD and Kenny by cleavage of DIAP2.

**Figure 4.**
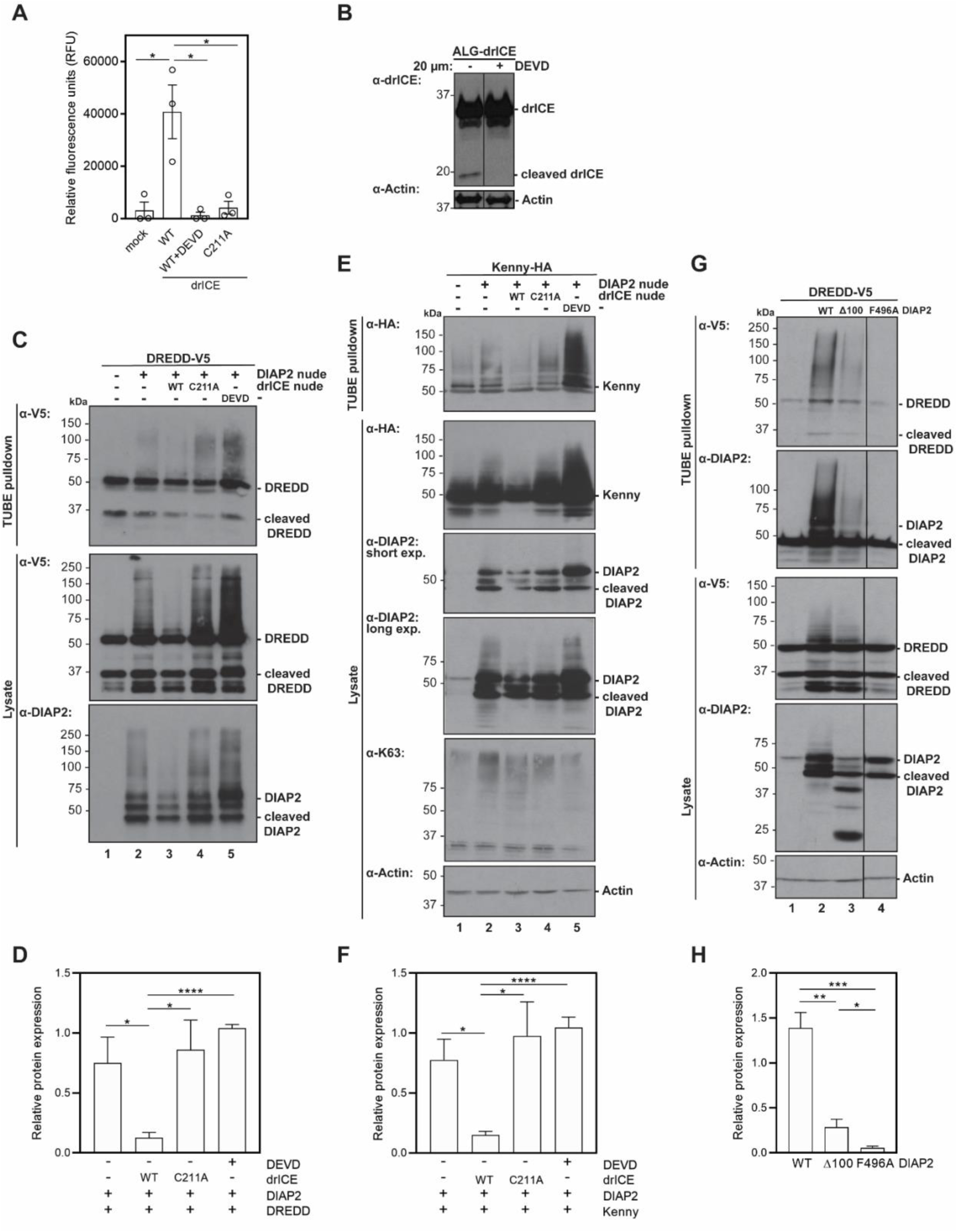
drICE interferes with the E3 ligase activity of DIAP2. A) *Drosophila* S2-cells were transfected with empty vector, drICE_WT_ or drICE_C211A_, where after the cells were treated with 20 µM Z-DEVD-FMK 16 h. The caspase-3/-7 activity was analysed by adding Apo-ONE reagent to plated cells and measuring fluorescence at 499/521 nm. Error bars indicate ± SEM from three experimental repeats, * p < 0.05. B) *Drosophila* S2-cells were transfected with ALG-drICE and treated with Z-DEVD-FMK 16 h. Cells were lysed and drICE-cleavage was analysed by Western blotting with α-drICE and α-Actin antibodies, n=3. C, E, G) *Drosophila* S2 cells were transfected with empty vector, DIAP2_WT_, drICE_WT_, drICE_C211A_, V5-tagged DREDD (C) or with HA-tagged Kenny (E), or with empty vector, V5-tagged-DREDD, DIAP2_WT_, DIAP2_Δ100_ and DIAP2_F496A_ (G), where after the cells were treated with 20 µM Z-DEVD-FMK for 16 h. Ubiquitin chains were isolated with GST-TUBE at denaturing conditions and the samples were analysed by Western blotting with α-V5, α-HA, α-DIAP2, α-K63 and α-Actin antibodies, n=3. D, F, H) The relative expression of ubiquitinated DREDD (D, H) or Kenny (F) was quantified. Error bars indicate ± SEM from at least three experimental repeats, * p < 0.05, ** p < 0.01, *** p < 0.001, **** p < 0.0001.

### drICE does not restrain pathogen induced inflammatory signalling

As drICE seem to restrain DIAP2’s ability to induce inflammatory signal transduction, the immune response upon septic pathogen-infection was tested in *drICE* overexpressing and in *drICE*-mutant flies. Interestingly, the effect of drICE is lost after septic infection with *Ecc15*, as *CantonS* control flies, *drICE*-mutant flies, *drICE-RNAi* flies, as well as flies overexpressing wild type *drICE* (**Figure 5A, B, C**) show a similar induction of *Drosocin* and *Diptericin* 5 h post infection. Furthermore, when monitoring the survival of flies, neither control flies, *drICE*_*17*_ flies nor *drICE* overexpressing flies, succumb to septic infection with *Ecc15* (**Figure 5D, E**). When studying the ability of DIAP2_Δ100_-expressing flies to mount a septic immune response, we found that these flies were both able to induce expression of *Drosocin* and *Diptericin* in response to infection **(Figure 5F**) and to survive septic infection similarly well as wild type *CantonS* and DIAP2_WT_ expressing flies (**Figure 5G**). These results indicate that the drICE-cleaved DIAP2_Δ100_ is able to mediate septic immune responses in the fat body. To investigate if the drICE-mediated regulation of DIAP2 was restricted to basal conditions, we analysed the intestinal immune responses to ingested pathogen. By performing a bacterial colony count after a 24-h feeding on *E. coli* and sucrose, we found that *diap2*_*7c*_;*diap2*_*Δ100*_-mutant (**Figure 5H**) and drICE overexpressing flies (**Figure 5I**) were able to fend off pathogens ingested with food similarly well as *diap2*_*7c*_;*diap2*_*WT*_ or control flies. This suggest that drICE only regulates basal immune responses, and not responses induced during pathogen infection.

**Figure 5.**
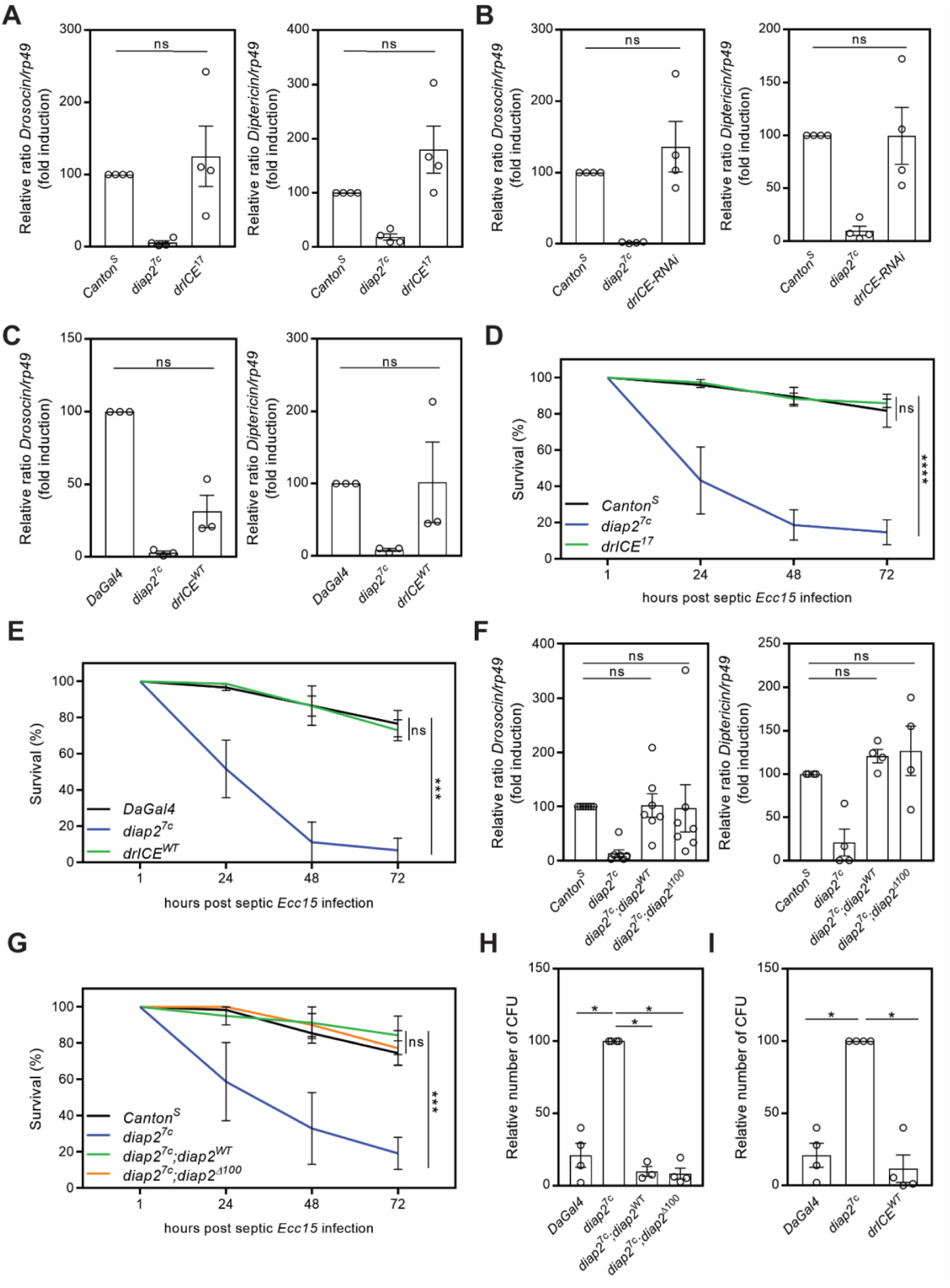
drICE does not restrain pathogen induced inflammatory signalling. A, B, C) Relative *Drosocin* and *Diptericin* mRNA levels analysed with qRT-PCR in *CantonS, diap2*_*7c*_, *drICE*_*17*_ (A) *UbiGal4;UAS-drICE-RNAi* (B) and in *DaGal4, diap2*_*7c*_ and *UAS-drICE*_*WT*_;*DaGal4* (C) flies 5 h after septic infection with the Gram-negative bacteria *Ecc15*. D, E) Adult *CantonS, diap2*_*7c*_ and *drICE*_*17*_ (D) or *DaGal4, diap2*_*7c*_ and *UAS-drICE*_*WT*_;*DaGal4* (E) flies were subjected to septic injury with *Ecc15* and their survival was monitored over time. F) Relative *Drosocin* and *Diptericin* mRNA levels analysed with qRT-PCR in adult *CantonS, diap2*_*7c*_, *diap2*_*7c*_;*UAS-diap2*_*WT*_*/DaGal4* and in *diap2*_*7c*_;*UAS-diap2*_*Δ100*_*/DaGal4* flies 5 h after septic infection with *Ecc15*. G) Adult *CantonS, diap2*_*7c*_, *diap2*_*7c*_;*UAS-diap2*_*WT*_*/DaGal4* and *diap2*_*7c*_;*UAS-diap2*_*Δ100*_*/DaGal4* flies were subjected to septic injury with *Ecc15* and their survival was monitored over time. H, I) *DaGal4, diap2*_*7c*_, *diap2*_*7c*_;*UAS-diap2*_*WT*_*/DaGal4* and *diap2*_*7c*_;*UAS-diap2*_*Δ100*_*/DaGal4* (H) or *UAS-drICE*_*WT*_;*DaGal4* (I) flies were infected by feeding with *E. coli* for 24 h and the bacterial load was assessed by counting colony-forming units (CFU). The same controls were used in H and I. Error bars indicate SEM from at least three independent experimental repeats, ns=non-significant, * p < 0.05, ** p < 0.01, *** p < 0.001, **** p < 0.0001.

### Receptor triggering by the commensal microbiome is needed to activate drICE

Finally, we wanted to investigate how and why both DIAP2 and drICE are activated in the intestinal epithelia. As the bacterial presence is constant in the gut, and the fat body only encounters bacteria during a systemic infection (Douglas, 2015), we hypothesized that pattern recognition receptors activate DIAP2-mediated IMD-signalling in the absence of drICE. To eliminate the commensal intestinal microbiome, we reared flies under axenic conditions. As expected, the expression of *Drosocin* and *Diptericin* was no longer elevated in axenic DIAP2_WT_ expressing flies (**Figure 6A**). Likewise, overexpression of DIAP2_WT_ in S2 fly cells could not induce AMP expression in the absence of receptor activation, here induced by co-expression of the receptor PGRP-LCx (**Figure 6B**). When rearing *UbiGal4*>*drICE RNAi* and *UbiGal4* driver flies under axenic conditions and measuring the expression of Relish target genes, we similarly found a significantly decreased expression of *Drosocin* and *Diptericin* in axenic *drICE RNAi* flies compared to conventionally reared *drICE RNAi* flies (**Figure 6C**). Furthermore, both wild type axenic flies and PGRP-LC mutant flies display lower caspase activity in the intestine compared to conventionally reared flies and control flies, respectively (**Figure 6D, E**), indicating a connection between the commensal bacteria in the gut, caspase activity and the ability of drICE to restrain IMD signalling. Taken together, we propose that commensal bacteria triggers formation of an initial PGRP-LCx receptor complex, aiming at recruiting DIAP2 to further activate the IMD pathway. The unnecessary inflammatory response is halted by drICE binding to, and forming an inhibitory complex with DIAP2, interfering with the ability of DIAP2 to induce downstream signalling and NF-κB target gene activation (**Figure 6F, G**).

**Figure 6.**
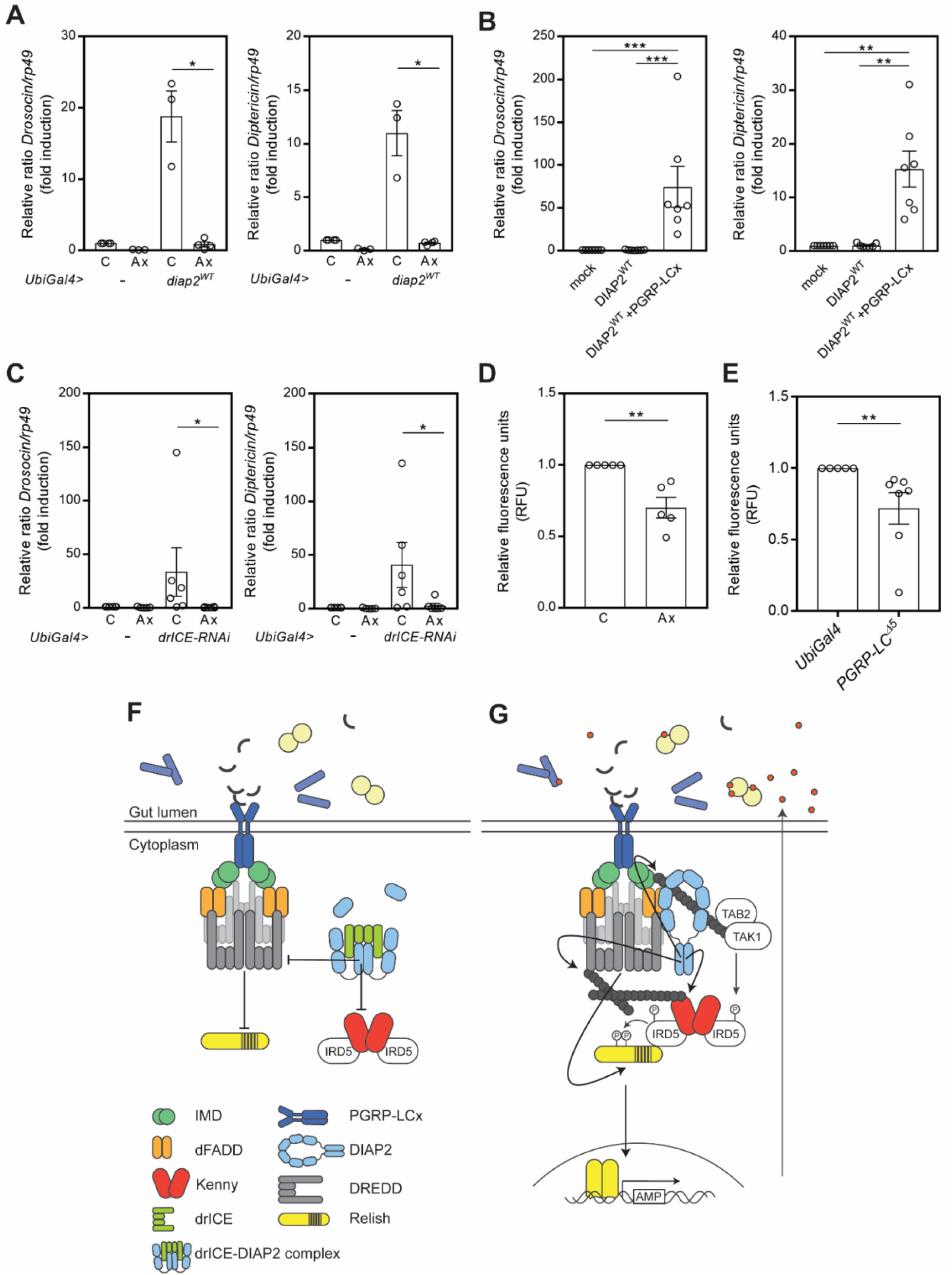
Receptor triggering by the commensal microbiome is needed to activate drICE. A) Relative *Drosocin* and *Diptericin* mRNA levels analysed with qRT-PCR in conventionally reared (C) or axenic (Ax) adult *UbiGal4* and *UbiGal4;UAS-diap2*_*WT*_ flies. Error bars indicate ±SEM from three independent experimental repeats, * p < 0.05. B) Relative *Drosocin* and *Diptericin* mRNA levels analysed with qRT-PCR from *Drosophila* S2 cells transfected with empty vector, DIAP2_WT_ and PGRP-LCx. Error bars indicate ±SEM from seven independent experimental repeats, ** p < 0.01, *** p < 0.001. C) Relative *Drosocin* and *Diptericin* mRNA levels analysed with qRT-PCR in conventionally reared (C) or axenic (Ax) adult *UbiGal4* and *UbiGal4;UAS-drICE-RNAi* flies. Error bars indicate ±SEM from six independent experimental repeats, * p < 0.05. D) Caspase-3/-7 activity in adult female guts of conventionally reared (C) or axenic (Ax) *yellow white* control flies was analysed by measuring fluorescence at 499/521 nm after addition of Apo-ONE reagent. Error bars indicate ±SEM from five independent experimental repeats, ** p < 0.01. E) Caspase-3/-7 activity in adult female guts of *PGRP-LC*_*Δ5*_ mutant flies and *UbiGal4* control flies was analysed by measuring fluorescence at 499/521 nm after addition of Apo-ONE reagent. Error bars indicate ±SEM from five independent experimental repeats, ** p < 0.01. F) PGN from the cell wall of commensal bacteria induce a low activation of the PGRP-LCx receptor, leading to the recruitment of IMD, dFADD and DREDD to the receptor complex. To avoid unnecessary NF-κB activation, active drICE binds to DIAP2, forming the inhibitory drICE-DIAP2 complex. drICE restrains DIAP2 from interacting with the pathway inducers DREDD and Kenny, bringing the IMD pathway to a stop. G) In the absence of drICE, DIAP2 is free to interact with, and ubiquitinate IMD, DREDD and Kenny. Ubiquitination of IMD enables recruitment of the TAB2-TAK1 complex, were after TAK1 can activate IRD5 by phosphorylation. IRD5 in turn phosphorylates Relish. Ubiquitination-dependent activation of DREDD and DREDD-meditated cleavage of Relish precedes translocation of the Relish dimer to the nucleus and subsequent activation of Relish target genes. Excessive amounts of AMPs, targeting commensal bacteria, are secreted to the gut lumen, leading to a disturbed gut homeostasis.

## Discussion

Caspases were first identified as regulators of apoptosis, but are now known to regulate other cellular processes such as immune signalling and development. Dysregulation of caspases has been implied in tumorigenesis, autoimmunity, autoinflammation and infectious pathologies (Van Opdenbosch and Lamkanfi, 2019, Halaby, 2012, Van Gorp et al., 2019). Caspases are known to be regulated by IAP proteins, however, regulation of the activity of IAP proteins by caspases has not been established before. Here, we report that the *Drosophila* caspase drICE functions as a negative regulator of intestinal immune responses regulated via the *Drosophila* NF-κB Relish. While drICE does not seem to impact pathogen-induced immune signalling, it restrains responses induced by commensal bacteria by interfering with the ability of DIAP2 to ubiquitinate its substrates DREDD and Kenny.

We suggest that the low activation of PGRP-LCx induced by commensal bacteria is sufficient to induce DIAP2-mediated Relish activation in the absence of a brake. In the intestinal epithelia, the brake is provided by the effector caspase drICE, halting unwanted inflammatory responses. Caspase-8-mediated cleavage of RIPK1 is essential to suppress inflammation in mammals and loss-of caspase-8 has been associated with immunodeficiency or early onset inflammatory bowel disease in humans (Newton, 2020, Lehle at al., 2019, Chun et al., 2002). We report, a similar caspase-mediated regulatory step, suppressing spontaneous inflammation in the form of drICE-mediated cleavage of DIAP2. Interestingly, flies expressing only the drICE-cleaved form of DIAP2, are able to induce inflammatory signalling in the gut in response to pathogen infection. This indicates that drICE is able to extend its regulatory effect post disruption of the DIAP2-drICE complex only during basal conditions. The constitutive interaction between DIAP2 and drICE in the intestinal epithelia may also be a consequence of a particular need to regulate caspase-activation to avoid apoptosis-induced cell proliferation (Fogarty and Bergmann, 2017).

The intestinal epithelial cells coexist with commensal bacteria and need to develop tolerance to pattern recognition to allow for healthy host-microbe interactions. However, these epithelial cells also need to maintain responsiveness to foodborne pathogens. When DIAP2 is cleaved and the drICE-DIAP2 complex is disassembled, also the BIR1 domain of DIAP2 is separated from the complex. As the cleaved DIAP2 functions as an IMD pathway-inducer during pathogen infection, separation of BIR1 seems to restrict intestinal immune signalling only during basal conditions. The DIAP2 homologue XIAP is a key regulator of NOD2-induced inflammatory signalling in the mammalian intestine, shown to induce the pathway by ubiquitinating RIPK2 (Hasegawa et al., 2008, Krieg et al., 2009). Interestingly, the nonsense mutation E99X in XIAP introducing a stop codon after the BIR1 domain, was found in an early onset Crohn’s disease patient. The mutation induced a severe and selective defect in intestinal NOD signalling, without affecting immune signalling in T cells and peripheral blood mono-nuclear cells (Zeissig et al., 2015). This suggests that a separated BIR1 domain is associated with impaired NF-κB signalling also in mammalian intestinal cells. The BIR1 domain has been shown to regulate XIAP-mediated NF-κB signalling, by bringing TAB1 and XIAP together, leading to activation of TAK1 (Lu et al., 2007). Hence, it will be interesting to study if the BIR1 of DIAP2 mediates NF-κB responses via, the yet unidentified, *Drosophila* TAB1-homologue in the epithelial cells of the fly intestine.

Proper regulation of NF-κB signalling is crucial, as de-regulated immune responses and chronic inflammation are deleterious for the host (Pasparakis, 2012). We propose that IAP-regulation by separating the BIR1 domain may provide a mechanism of restraining unwanted inflammatory responses induced in the cells of the epithelial tissues exposed to non-pathogenic microbiota. While IAP cleavage does not seem to affect immune cell-mediated pathogen-induced responses, this may allow for specific regulation of malfunctional epithelial responses in chronic inflammatory disease.

## Material and Methods

### Fly husbandry and strains

*Drosophila melanogaster* were maintained at 25°C with a 12 h light-dark cycle on Nutri-fly BF (Dutscher Scientific). The *CantonS* strain was used as wild type flies. *DaGal4* and *NP1-Gal4* driver lines, *Diptericin-LacZ* reporter lines, and *diap2*_*7c*_ mutants and transgenes were provided by Dr. François Leulier. The *UbiGal4* strain was provided by Dr. Ville Hietakangas, and *drICE*_*17*_ mutants by Prof. Andreas Bergmann. *PGRP-LCΔ5* (stock #36323) and *UAS-p35* (stock #5073) strains were obtained from Bloomington *Drosophila* stock centre and *drICE-IR* (stock #28064) flies were from the Vienna *Drosophila* Resource Centre. V5-tagged *drICE*_*WT*_ and *drICE*_*C211A*_, as well as untagged *DIAP2*_*WT*_ and *DIAP2*_*Δ100*_ were cloned into pUAST, and transgenic flies were generated by BestGene inc. Axenic flies were reared germ-free according to the previously published protocol (Kietz et al., 2018). In short, fly embryos were de-chorionated using 2% active hypochlorite, and washed twice in 70% ethanol and sterile H_2_O. After removal of the chorion, eggs were placed in autoclaved food and left to develop in a sterile environment. The hatched flies were confirmed to be axenic by 16S PCR and by growing fly homogenates on Luria Bertani (LB) plates and checking for bacterial growth.

### Bacterial strains, infection and survival experiments

The Gram-negative bacteria *Erwinia carotovora carotovora 15* (*Ecc15*) was kindly provided by Dr. François Leulier and the *E. coli* Top10 strain was purchased from Thermo-Fisher Scientific. *Ecc15* used for septic infection was cultivated in LB media at 29°C for 16-18 h on agitation and concentrated (optical density of 200). To induce septic injury, adult flies were pricked in the lateral thorax with a needle previously dipped in concentrated *Ecc15* solution. For quantitative PCR (qPCR) 10 adult flies were incubated 5 h at 25°C after infection. For survival assays at least 20 adult flies were counted at indicated time points after infection. Infection experiments were excluded if more than 25% of the negative control strains survived bacterial infection or if AMP gene expression was significantly enhanced in these flies. In these cases, the bacterial potency was considered too low. Survival experiments in which wild-type flies survived to a lesser extent than 75%, were also excluded. These criteria were pre-established. For bacterial colony count, *Escherichia coli* (*E. coli*) was transformed with pMT/Flag-His-amp and cultivated in ampicillin-containing LB medium at 37°C for 16–18 h on agitation and concentrated by centrifugation (optical density of 100). After a 2 h starvation, adult flies were fed for 24 h with a 1:1 solution of transformed *E. coli* in 5% sucrose at 25°C. Four flies were cleaned with ethanol and sterile H_2_O, and homogenised in 300 µl PBS. Samples were diluted 1:1000 and plated on LB agar plates containing 50 µg/ml ampicillin. Colonies were counted 24 h after plating.

### Cell culture and transfection of *Drosophila* S2 cells

*Drosophila* Schneider S2 cells (Invitrogen) were grown at 25°C using Schneider’s cell medium supplemented with 10% fetal bovine serum, 1% L-glutamine and 0.5% penicillin/streptomycin. S2 cells were transfected with indicated constructs using Effectene transfection reagent (QIAGEN) according to the manufacturer’s instructions. For lysis with Laemmli sample buffer and for GST-pulldown assays, respectively 2 x 10^6^ and 0.7 x 10^7^ cells were seeded prior to transfection. Expression of pMT plasmids was induced with 500 µM CuSO_4_ 16 h before lysis. The caspase inhibitor Z-DEVD-FMK (BD Pharmingen) was used at 20 µM 16 h before lysis.

### Plasmids and antibodies

The plasmids pMT/V5His and pAc5/V5His (Invitrogen) were used as backbones for tag insertions and removals, and for subcloning of the constructs pMT/FlagHis, pMT/HAFlag, pAc5/DIAP2, pMT/DREDD-V5His, pMT/PGRP-LCx-V5His, pMT/Kenny-HA, pMT/drICE, pMT/ALG(p20/p10)drICE-V5Flag and pAc5/DIAP2_Δ100_. Point mutations DIAP2_F496A_ and drICE_C211A_ were made by site-directed mutagenesis (Agilent Technologies or Stratagene). The following antibodies were used: α-K63 (clone Apu3, #05-1308, Millipore), α-DIAP2 (Tenev et al., 2005), α-drICE (Leulier et al., 2006b), α-HA (clone 3F10, #11867423001, Roche), α-V5 (Clone SV5-Pk1, #MCA1360, Bio-Rad), α-phospho-histone H3 (PH-3) (Ser10, #9701, Cell Signalling Technology) and α-Actin (C-11, sc-1615, Santa Cruz).

### Purification of GST-TUBE-fusion protein

GST-TUBE expression was induced in *E. coli* BL21 by adding 0.5 mM IPTG to an overnight culture of bacteria in LB medium at 18°C. Bacteria were lysed by sonication in lysis buffer containing PBS, 300 mM NaCl, 1mM DTT, and protease inhibitor Complete EDTA-free (Roche). The lysate was incubated with Gluthatione Sepharose™ 4B (GE Healthcare) for 2 h and then washed in wash buffer containing 200 mM Tris (pH 8) and 1.5 M NaCl. GST-TUBE was eluted in 50 mM Tris (pH 8.5), 150 mM NaCl, 10% Glycerol, 2 mM β-mercaptoethanol and 20 mM gluthatione.

### Purification of ubiquitin conjugates from cells

Ubiquitin conjugates were purified using a recombinant protein containing four ubiquitin binding entities in tandem (tandem ubiquitin binding entity, TUBE) fused to GST (GST-TUBE). Cells were lysed in a buffer containing 20 mM NaH_2_PO_4_, 20 mM Na_2_HPO_4_ 1% NP-40 and 2 mM EDTA supplemented with 1 mM DTT, 5 mM NEM, Pierce™ Protease and Phosphatase Inhibitor, 5 mM chloroacetamide and 1% SDS. Lysates were sonicated, diluted to 0.1% SDS, and cleared before incubation with Glutathione Sepharose™ 4B (GE Healthcare) and GST-TUBE for a minimum of 2 h under rotation at 4°C. The beads were washed four times with ice cold PBS-0.1% Tween-20 and eluted using Laemmli sample buffer.

### Lysis of S2 cells and *Drosophila* whole flies or organs for Western blotting

Transfected S2 cells were harvested in ice cold PBS and lysed in Laemmli sample buffer. Adult *Drosophila* flies, or dissected intestines or abdomens from adult female flies, were homogenized and lysed 10 min on ice in a buffer containing 50 mM Tris (pH 7.5), 150 mM NaCl, 1% Triton X-100, 1 mM EDTA and 10% Glycerol. The lysates were cleared before addition of Laemmli sample buffer.

### Quantitative RT-PCR (qPCR)

*Drosophila* adult flies or *Drosophila* S2 cells were homogenised using QIAshredder (QIAGEN) and total RNA was extracted with RNeasy Mini Kit (QIAGEN) according to the manufacturer’s protocol. cDNA was synthesised with iScript cDNA synthesis kit (Bio-Rad) according to the manufacturer’s protocol. qPCR was performed using SensiFAST™ SYBR Hi-ROX Kit (Bioline). *rp49* was used as a housekeeping gene for ΔCt calculations. The following gene specific primers were used to amplify cDNA: *Diptericin* (5’-ACCGCAGTACCCACTCAATC, 5’-ACTTTCCAGCTCGGTTCTGA), *Drosocin* (5’-CGTTTTCCTGCTGCTTGC, 5’-GGCAGCTTGAGTCAGGTGAT), *rp49* (5’-GACGCTTCAAGGGACAGTATCTG, 5’-AAACGCGGTTCTGCATGAG).

### Immunofluorescence of *Drosophila* intestines

Intestines from female adult flies were dissected in PBS and fixed 10 minutes in 4% paraformaldehyde. Samples were permeabilised with PBS-0.1% Triton X-100, 1 h at room temperature, washed with PBS and incubated over night at 4°C with primary antibody PH-3 (Ser10, #9701, Cell Signalling Technology) 1:1000. After washing, the intestines were incubated 2 h at room temperature with secondary antibody Alexa Fluor 488 donkey anti-rabbit IgG (#A21206, Invitrogen) 1:600. Both primary and secondary antibodies were diluted in PBS and 1% bovine serum albumin. DNA was stained with DAPI (4’,6-diamidino-2-phenylindole) (Invitrogen). After washing with PBS, the samples were mounted using Mowiol (Sigma). Imaging was performed with a spinning disk confocal microscope (Zeiss Axiovert-200M microscope Yokogawa CSU22 spinning disk confocal unit) using x20 objectives. The 3i SlideBook6 software was used for image acquisition and image processing was done with Image J.

### X-gal staining of *Drosophila* intestines and abdomens

Intestines or abdomens from adult female flies were dissected in PBS and fixed 15 minutes at room temperature with PBS containing 0.4% glutaraldehyde and 1 mM MgCl_2._ The samples were washed with PBS and incubated with fresh staining solution containing 5 mg/ml X-gal, 5 mM potassium ferrocyanide trihydrate, 5 mM potassium ferrocyanide crystalline and 2 mg/ml MgCl_2_ in PBS 1 h at 37°C. After washing with PBS, the samples were mounted in Mowiol (Sigma) and imaged with bright field microscopy (Leica).

### Fluorometric measurement of caspase-3/7 activity in cells or in fly intestines or abdomens

Transfected S2 cells were plated in a 96-well plate or intestines or abdomens from adult female flies were dissected in PBS and lysed in buffer containing 50 mM Tris (pH 7.5), 150 mM NaCl, 1% Triton X-100, 10% glycerol, 1 mM EDTA and Pierce™ Protease Inhibitor. The lysate was cleared at 12 000 rpm for 10 min at 4°C and protein concentration adjusted with Bradford assay (BioRad). The caspase-3/7 activity of the lysates or the plated cells was analysed using Apo-ONE ® Homogenous Caspase-3/7 Assay (Promega) according to manufacturer’s protocol. Fluorescence was measured at 499/521 with the plate reader HIDEX sense (HIDEX).

### Sequencing of the 16S rRNA gene

Genomic DNA was isolated from 40 adult flies using a modified protocol for the QIAamp DNA mini kit (QIAGEN) (Simhadri et al., 2007). Flies were surface sterilized by vortexing them twice in 2% active hypochlorite and sterile H_2_O. The efficiency of the washes was confirmed by 16S PCR of water from the last wash step. Flies were homogenized in lysis buffer containing 20 mM Tris, pH 8.0, 2 mM EDTA, 1.2% Triton X-100 and 20mg/ml lysozyme, and incubated 90 min at 37°C. 200 µl AL buffer (QIAamp DNA mini kit) with 20 µl proteinase K were added and the lysate was incubated 90 min at 56°C. Subsequent extraction was performed according to manufacturer’s protocol. Amplification and Illumina MiSeq sequencing of the V1-V3 region of the 16S rRNA gene, as well as selection of operational taxonomic units (OTUs) and taxonomy assignment of OTUs was done using Eurofins Genomics InView Microbiome Profiling 3.0 service. The sequenced *CantonS* flies were positive for *Wolbachia* species. The proportion of *Wolbachia* species have been omitted in Figure 2I for easier comparison of bacterial species residing in the gut lumen.

### Statistical analysis

Results from survival assays were analysed by two-way analysis of variance (ANOVA) with Tukey’s post hoc test for 95% confidence intervals and results from qPCR by Student’s t-test on the ΔΔCt value, graphs depict relative fold induction of the target gene compared to a normalised control sample. In comparison to normalised control values the Mann-Whitney U test was applied. Relative protein expression from Western blots were measured with ImageJ and analysed by Student’s t-test. In figures ns stands for p > 0.05, * for p < 0.05, ** for p < 0.01, *** for p < 0.001 and **** for p < 0.0001. Error bars in figures specify ±SEM from the indicated number of independent experiments. The experiments were repeated at least three times. With smaller differences in detection, more repeats were done.

## Acknowledgements

The HIDEX plate reader was provided through collaborator with Prof. Kid Törnquist. We are grateful to Anna Ahlbäck and Dr. Tencho Tenev for technical support and Francois Leulier for fly strains, plasmids and bacterial strains as mentioned. The Academy of Finland (#275570, #283524, #312557), the Magnus Ehrnrooth Foundation, the Sigrid Jusélius Foundation, The Foundation Liv och Hälsa, the Åbo Akademi University Foundation, Instrumentarium Science Foundation and Turku Doctoral Network in Molecular Biosciences are acknowledged for financial support.

## Author Contributions

CK designed and executed most of the experiments, data analysis and writing of the manuscript. VP planned and performed axenic fly experiments, I-ET, PR and AM performed survival and AMP analyses of DIAP2 and drICE mutants. PR, PM and AM contributed to the design of the experiments, writing and data analysis of this manuscript.

